# Ancestral chromatin state constrains the functional landscape of bivalent domains in mammalian spermatogenesis

**DOI:** 10.64898/2026.04.14.718508

**Authors:** Delaney Farris, Justin Tai, Bluma Lesch

## Abstract

Mammalian germ cells are enriched for bivalent chromatin, an epigenetic state defined by the dual presence of the activating H3K4me3 and repressive H3K27me3 histone modifications. Bivalency is evolutionarily conserved at developmentally important genes in germ cells but diverges at hundreds of additional loci, and evolutionary gains in bivalency have been proposed to reflect divergent somatic functions of the associated genes. Here, we sought to discover if evolutionary gains in bivalency occur selectively at genes with specific functions, and to better elucidate the role of bivalent chromatin in germ cells. By comparing genome-wide profiles for four histone modifications in spermatogenic cells of six mammalian species, we define a comprehensive set of mammalian bivalent domains and classify them based on conservation or divergence of chromatin state. We find that evolutionarily conserved bivalent regions exhibit canonical features of bivalency and maintain bivalency in embryonic stem cells. In contrast, bivalent domains emerging from a purely active or repressed ancestral chromatin state have atypical sequence and regulatory features and are frequently germ cell specific. Genes associated with these recent bivalent domains exhibit distinct somatic expression patterns that reflect their ancestral chromatin state in germ cells. Specifically, bivalent genes emerging from ancestrally active chromatin are more highly expressed in somatic tissues and are enriched for immune-related functions, while those emerging from ancestrally H3K27me3-only domains are lowly expressed in the soma and enriched for neurogenesis functions. We propose that recent bivalent regions demarcate sites of regulatory sequence change that preferentially impacts specific somatic lineages.

## Introduction

Bivalent chromatin is an epigenetic regulatory state defined by the dual presence of the opposing histone modifications H3K27me3 and H3K4me3 (trimethylation of histone H3, lysines 27 and 4, respectively) and whose biological meaning remains elusive. Bivalency was initially characterized in embryonic stem cells (ESCs)^1–3^ and is found at promoters of developmental genes in mammalian cell types with pluripotent potential, including primordial germ cells^4,5^ and preimplantation embryos^6^, and at lower levels in some more differentiated cellular contexts^7–9^. Disruption of bivalency-associated factors such as MLL2/KMT2B^10^, PRC2^11,12^ and PRC1^13^ in ESCs results in deregulation of bivalent gene expression and subsequent impairment of differentiation, leading to the hypothesis that bivalency regulates differentiation potential. However, despite its likely significance in guiding differentiation and cell fate, the specific in vivo biological functions of bivalency and mechanisms that establish cell type-specific bivalency patterns have not been resolved.

Bivalency is abundant in mature male and female mammalian germ cells^14–17^. In male germ cells, bivalency is present throughout development, and a subset of genes newly gains bivalency at the meiotic stage of spermatogenesis^18,19^. Likewise, during mouse oocyte maturation, there is a dramatic increase in regions marked with bivalent chromatin.^17^ The unique dynamics and regulatory machinery of bivalency in germ cells suggest that its establishment and function in the germ line may differ from embryonic or somatic stem cells. Consistent with a distinct function, bivalency in germ cells has been suggested to have dual roles where it contributes both to maintaining germ cell identity and to generating a totipotent zygote^4,5,19^.

Bivalency is well conserved in male germ cells in mammals^14^, but a subset of regions gain or lose bivalency over evolutionary time. Gain and loss of bivalency has been suggested to parallel changing developmental function of associated genes^14^. Comparing the function and sequence of orthologous loci that differ in their bivalent status across species offers an opportunity to discover features that specify and regulate the bivalent chromatin state, similar to other comparative approaches that have identified patterns of regulatory divergence in testis^20,21^ and in diverse somatic tissues^22–26^. Interestingly, analysis of the evolutionary dynamics of gene expression in mammalian testes has led to the hypothesis that male germ cells are a “testing ground” for gene regulation and expression, where new regulatory states first appear in germ cells and are then reinforced in somatic tissues over many generations ^20,27–29^. Importantly, staged male germ cells isolated from adult testis are developmentally and biologically comparable across mammalian species and thus constitute an ideal system for comparing regulatory states, in contrast to ESCs, which require species-specific culture conditions and recapitulate different stages of preimplantation development^30^.

Here, we uncover distinct features of emerging bivalent domains by comparing the bivalent state in male germ cells of six mammalian species and in human and mouse ESCs. Applying a comprehensive genome-wide and multispecies approach, incorporating four histone modifications and comparing across germ cells and ESCs, we identify thousands of bivalent loci not previously characterized by past studies, including both gene-distal and gene-proximal regions, sites where bivalency is restricted to late spermatogenic cells and not present in ESCs, and regions with evolutionarily stable or dynamic bivalency ^3,14,19^. We find that bivalent promoters exhibit distinct sequence features and functions depending on their ancestral epigenetic status in germ cells. Evolutionarily conserved bivalent regions in germ cells often retain bivalency in ESCs, exhibit typical sequence features associated with bivalency, and regulate core developmental processes. In contrast, evolutionarily new bivalent domains in both primate and rodent lineages are less likely to be bivalent in ESCs and have distinct sequence features reflecting their evolutionary trajectory. Further, their transcriptional status and function in somatic tissue often reflects their ancestral, non-bivalent germline chromatin state, especially in neural and immune tissues. Our results suggest a germ cell-specific function for bivalency in buffering regulatory evolution in germ cells.

## Results

### Bivalent chromatin is abundant in mammalian male germ cells

To comprehensively map bivalency in germ cells across the mammalian lineage, we generated genome-wide chromatin profiles using new and public histone modification ChIP-seq data from male germ cells in six mammalian species: human, rhesus, mouse, rat, bull, and opossum ^14,31–34^, **Fig. 1A**, **Table S1**). Quality of all germ cell datasets was confirmed based on alignment rate, de-duplicated sequencing depth, and correlation between biological replicates (**Fig. S1**, **Table S2**). We applied ChromHMM (^35,36^) to define combinatorial chromatin states from the histone modifications H3K27me3, H3K4me3, H3K4me1, and H3K27ac independent of gene annotations. We chose a 12-state model that recovered similar states in all six species and annotated each state as one of four functionally relevant categories: Bivalent, Polycomb, ActiveTSS, and Enhancer-like (**Fig. 1B-C**, **Fig. S2A**). The bivalent label was assigned based on dual enrichment of H3K27me3 and H3K4me3. Interestingly, all species also showed enrichment for H3K4me1 in at least one ChromHMM state annotated as bivalent using this criterion, reinforcing H3K4me1 enrichment as an additional feature of bivalency ^31^. Quantification of H3K4me3 and H3K27me3 signal in all annotated regions showed that, as expected, ChromHMM states annotated as bivalent generally had high H3K4me3 and H3K27me3 signal, activeTSS regions had high H3K4me3 and lower H3K27me3 signal, and Polycomb regions had high H3K27me3 and lower H3K4me3 signal, confirming that the ChromHMM segmentation captures meaningful chromatin states (**Fig. 1D**).

**Figure 1:**
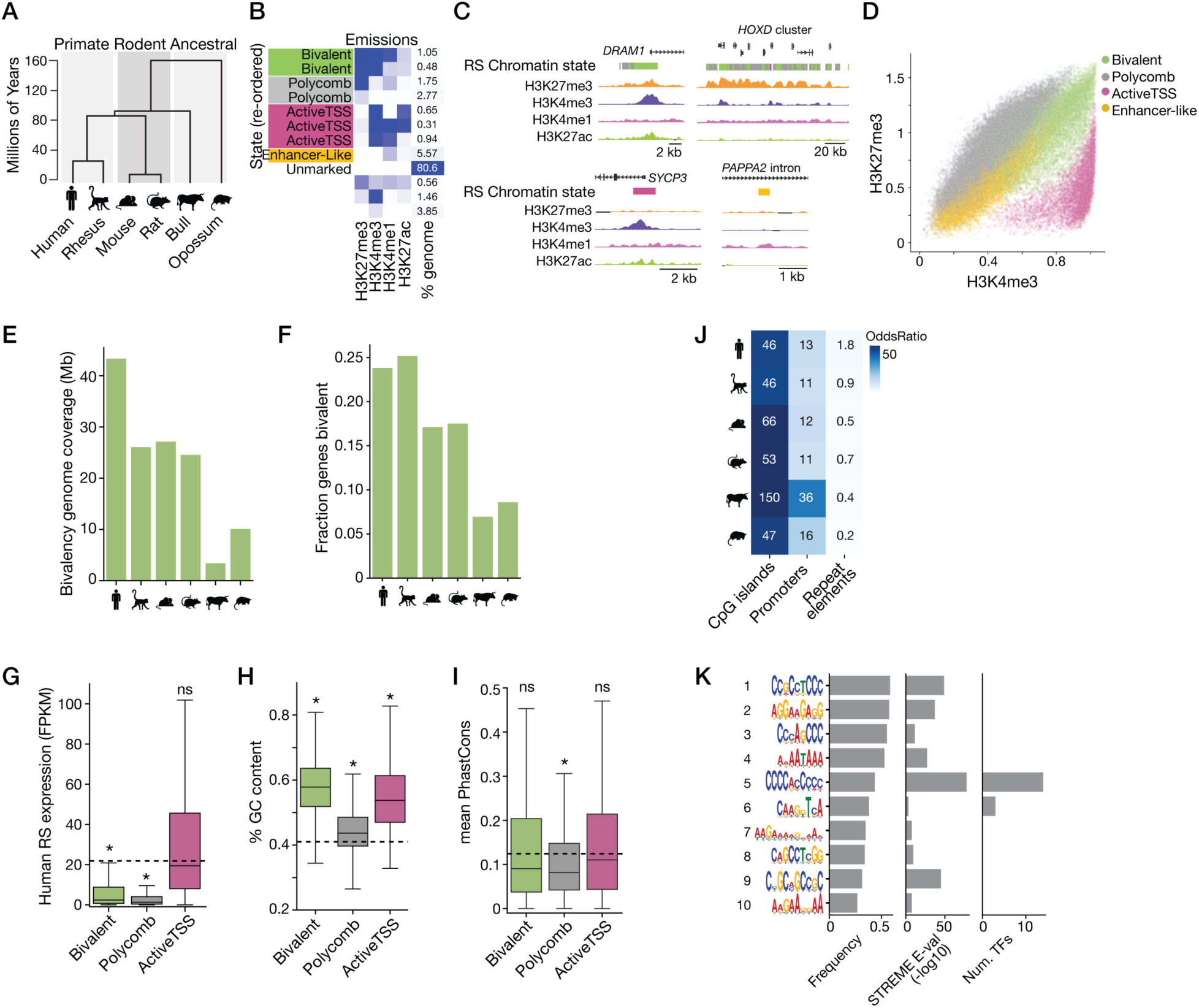
Characterization of bivalent chromatin across 6 mammalian species in the male germ line. **A,** Phylogeny of the species used in this study. **B,** ChromHMM emissions plot and genomic coverage for human round spermatid (RS) chromatin state maps. **C,** Example genome browser tracks (hg38) of chromatin states defined by the ChromHMM model. **D,** Quantitative H3K4me3 and H3K27me3 ChIP-seq signal information for individual regions defined by ChromHMM (see Methods). Color represents functional annotation (FA) of the ChromHMM state. **E,** Total genomic coverage in megabases (Mb) of bivalent regions across species. **F,** Fraction of promoters (+/- 1kb around TSS) which overlap with a bivalent domain. Total numbers of promoters: human n=86,363, rhesus n=35,419, mouse n=78,296, rat n=32,862, bull n=36,068, opossum n=30,367. **G,** Expression level (FPKM) of genes with promoters marked by a given chromatin state in human RS. Box shows median and interquartile range; whiskers show range of the distribution. *p < 1e-10, 1-sample t-test performed (alternative=”less”) against the mean expression of all genes (dashed line). **H,** GC content of individual regions within each FA type. *p < 1e-30, 1-sample t-test performed (alternative=”greater”) against the mean whole-genome GC content (hg38, dashed line). **I,** Mean phastCons score within individual regions by FA type. *p < 1e-30, 1-sample t-test performed (alternative=”less”) against the mean of a set of re-shuffled regions in the human genome (hg38, dashed line). **J,** Genomic enrichment (odds ratio) of CpG islands, promoters, and repeat elements (LINE/SINE/LTR) in bivalent elements across species (see Methods). **K,** Position weight matrix of the ten most frequent enriched DNA motifs found in human bivalent regions near promoters (n=10,000). For each enriched de-novo motif, frequency in the test set (left), motif E-value (-log10) (middle), and number of predicted TFs (right) are shown.

Bivalent chromatin segments covered approximately 43 Mb (0.01%) of the human genome, approximately 25 Mb (0.008-0.01%) of the rhesus, rat, and mouse genomes, and 3.4-10 Mb (0.001% and 0.003%) of the bull and opossum genome, respectively (**Fig. 1E**). Approximately 23% and 18% of promoters overlapped bivalent regions in primate and rodent species, respectively, and this number was 6-8% in bull and opossum, proportional to the total bivalency coverage in each species (**Fig. 1F**). Consistent with expectations, expression of genes with bivalent or Polycomb promoters was low in germ cells compared to the mean expression of all genes (p < 1e-30, 1-sample t-test), while expression of genes marked by activeTSS was comparable to the genome average (**Fig. 1G**). Also consistent with studies in ESCs, bivalent regions had an average GC content of 56%, slightly higher than active elements (54%) and substantially higher than the genome average (p < 1e-30, 1-sample t-test), or Polycomb regions (44%) (**Fig. 1H**). Overall levels of sequence conservation were similar among the functional annotation classes (**Fig. 1I**). As observed for bivalent regions defined in ESCs, bivalent segments were strongly enriched at CpG islands (>45-fold over expected), moderately enriched at promoters (>10-fold) and weakly enriched or depleted at repeat elements (LINEs, SINEs, and LTRs) in all six species (**Fig. 1J**).

We next sought to identify DNA sequence motifs enriched in human bivalent regions and the transcription factors (TFs) predicted to bind these motifs (Methods). Six of the ten most frequent motifs identified by STREME (^37,38^) (found in 31-58% of bivalent loci) consist mostly of repetitive C and G nucleotides (**Fig. 1K**, motifs 1-3, 5, 8-9) and are slightly longer than typical TF binding motifs. Only two of the ten most frequent enriched motifs, found in 44% and 38% of bivalent sequences, were predicted to be bound by a specific TF, including KLF/SP family TFs, ESRRA/B, MAZ, ZNF740, and NR6A1 (**Fig. S2B**). A parallel analysis of enriched motifs in mouse bivalent regions found very similar patterns and nearly identical predicted TFs (**Fig. S2B**, **S2C**). Expression of most of these predicted TFs is low in testis across species (**Fig. S2D**). KLF and SP family TFs commonly bind developmentally regulated genes, suggesting that enrichment of these motifs in bivalent regions likely reflects a correlation due to mutual enrichment at developmental gene promoters, rather than a role for these TFs in bivalency establishment or regulation, consistent with previous findings^39^. This result suggests that a simple TF-dependent mechanism is probably not sufficient to specify the bivalent state in germ cells.

### Bivalent chromatin is found at candidate regulatory elements both proximal and distal to promoters

An advantage of our unbiased genome segmentation approach is the ability to identify intergenic and other non-promoter bivalent elements. Indeed, we found that about half of the identified bivalent regions in each species were distal to promoters (> 1 kb from a transcriptional start site) (**Fig. 2A**). On average, promoter-distal bivalent regions were slightly shorter than promoter-proximal regions, with average lengths of 1,140 and 1,735 base pairs respectively (**Fig. 2B**). Promoter-distal bivalent regions also had lower overall H3K27me3 and H3K4me3 ChIP-seq signal compared to promoter-proximal bivalent regions (**Fig. 2C**), which could be explained in part by the smaller region size. However, promoter-distal bivalent regions had robust signal for both H3K27me3 and H3K4me3 as visualized on the genome browser (**Fig. 2D**). To assess if these distal bivalent regions could have regulatory function, we asked if they overlapped with known regulatory elements, specifically ENCODE-defined distal enhancer-like regions (dELS) and open chromatin sites (DNase+H3K4me3). We found that 3,766 and 2,466 human and mouse distal bivalent elements overlap with dELS, respectively, and 882 and 762 human and mouse distal bivalent elements overlap with open chromatin sites (**Fig. 2E**). The co-occurrence of bivalent regions with these regulatory elements is more than expected by chance (p-val < 1e-30, Fisher’s exact test). This finding suggests that promoter-distal bivalent regions could represent weak bivalent sites that mark latent genome regulatory elements, particularly enhancer-like regions. Together, these results indicate that bivalency is widespread in mammalian male germ cells, including in intergenic regions, and that germline bivalent regions overall largely recapitulate well-established features of bivalency previously defined in ESCs.

**Figure 2:**
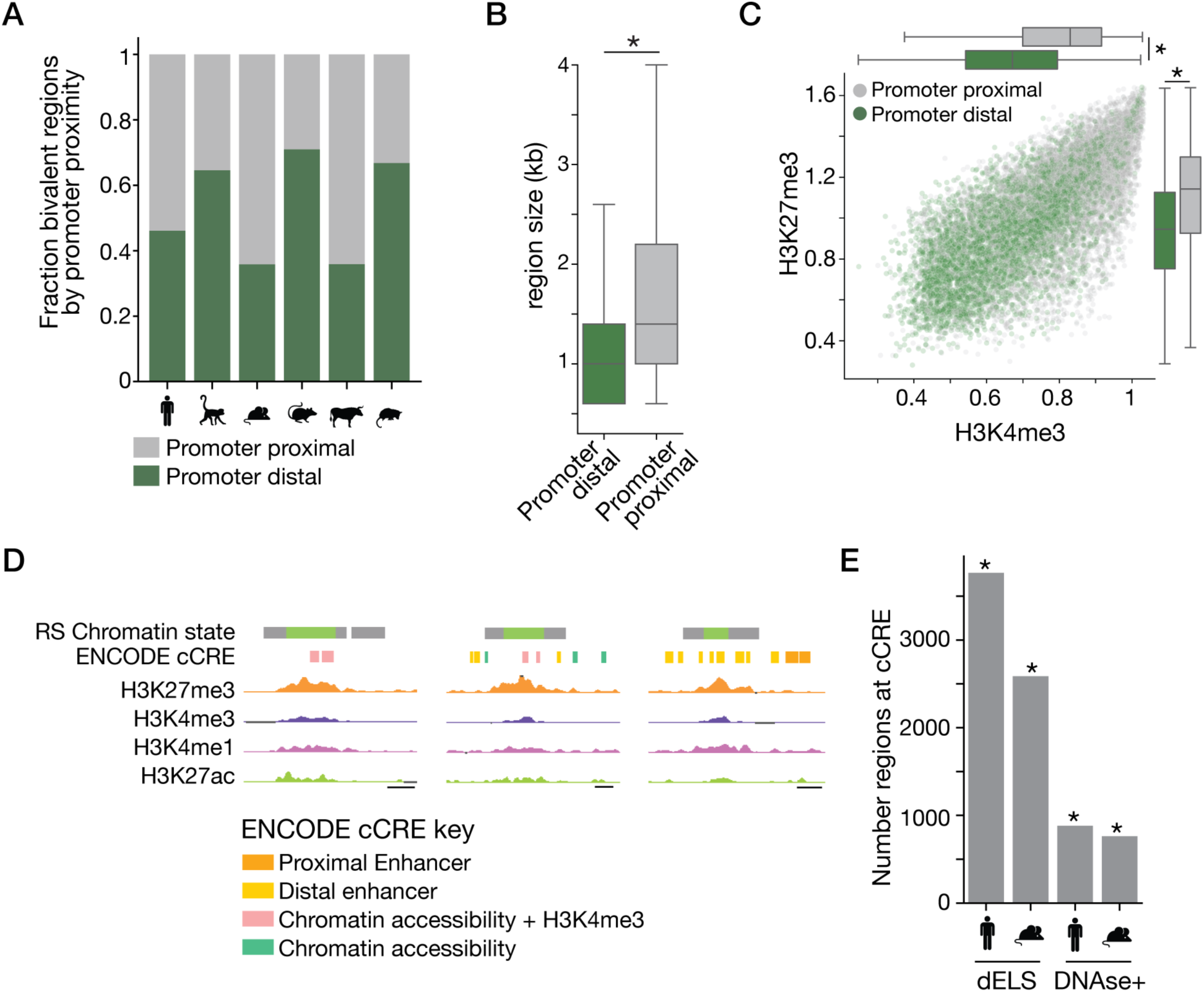
Bivalent chromatin exists outside of gene promoter proximal regions. **A,** Bivalent regions classified as promoter proximal (within +/- 1kb of all TSS) or promoter distal across species. Total numbers of bivalent elements considered: human n=28,478, rhesus n=17,828, mouse n=18,327, rat n=18,626, bull n=2,673, opossum n=8,094. **B,** Size of human bivalent regions classified by promoter proximity status. Box shows median and interquartile range; whiskers show range of the distribution. *p < 1e-30, Two-sample, two-sided Kolmogorov-Smirnov test performed between the two groups. **C,** Quantitative H3K4me3 and H3K27me3 ChIP-seq signal information for individual human bivalent regions stratified by promoter proximity. *p < 1e-30, two-sample, two-sided Kolmogorov-Smirnov test performed between the two groups. **D,** Example genome browser tracks of promoter-distal human bivalent regions. Scale bar equals 1 kb. **E,** Number of promoter-distal bivalent regions (human n=13,136, mouse n=6,575) that overlap with distal enhancer like elements (dELS) and open, active chromatin elements (DNase+H3K4me3) defined by ENCODE. *p < 1e-30, Fisher’s exact test performed to compute likelihood of co-occurrence of the two region sets.

### A large class of bivalent regions is specific to spermatogenic cells

Features usually considered typical of bivalency were defined based on data from ESCs. Since we found that bivalent regions in the male germ line overall share these features, we asked if the bivalent regions we identified in germ cells coincided with bivalent loci defined in ESCs. Using published data^40–42^, we generated analogous chromatin state maps based on ChromHMM segmentation in human and mouse ESCs (**Fig. 3A**, **Fig. S3A**, **Table S1, Table S2**). This segmentation recaptured over 85% of previously published high-confidence ESC bivalent regions in both species (**Fig. S3B**) ^10,43^. We compared the identified ESC bivalent regions to germline bivalent regions and found that 80% (human) and 70% (mouse) of bivalent regions in ESCs were also bivalent in the germ line. Surprisingly however, only 12% and 35% of germline bivalent regions were also bivalent in human and mouse ESCs, respectively, reflecting a much larger total set of bivalent regions in germ cells (**Fig. 3B**). This result is not due to many smaller regions being called as bivalent in germ cells since it was also reflected in total genomic coverage, where bivalency covers 2-6x more of the genome in germ cells compared to ESCs (**Fig. 3C**). We conclude that the bivalent state is significantly more prevalent in male germ cells compared to ESCs.

**Figure 3:**
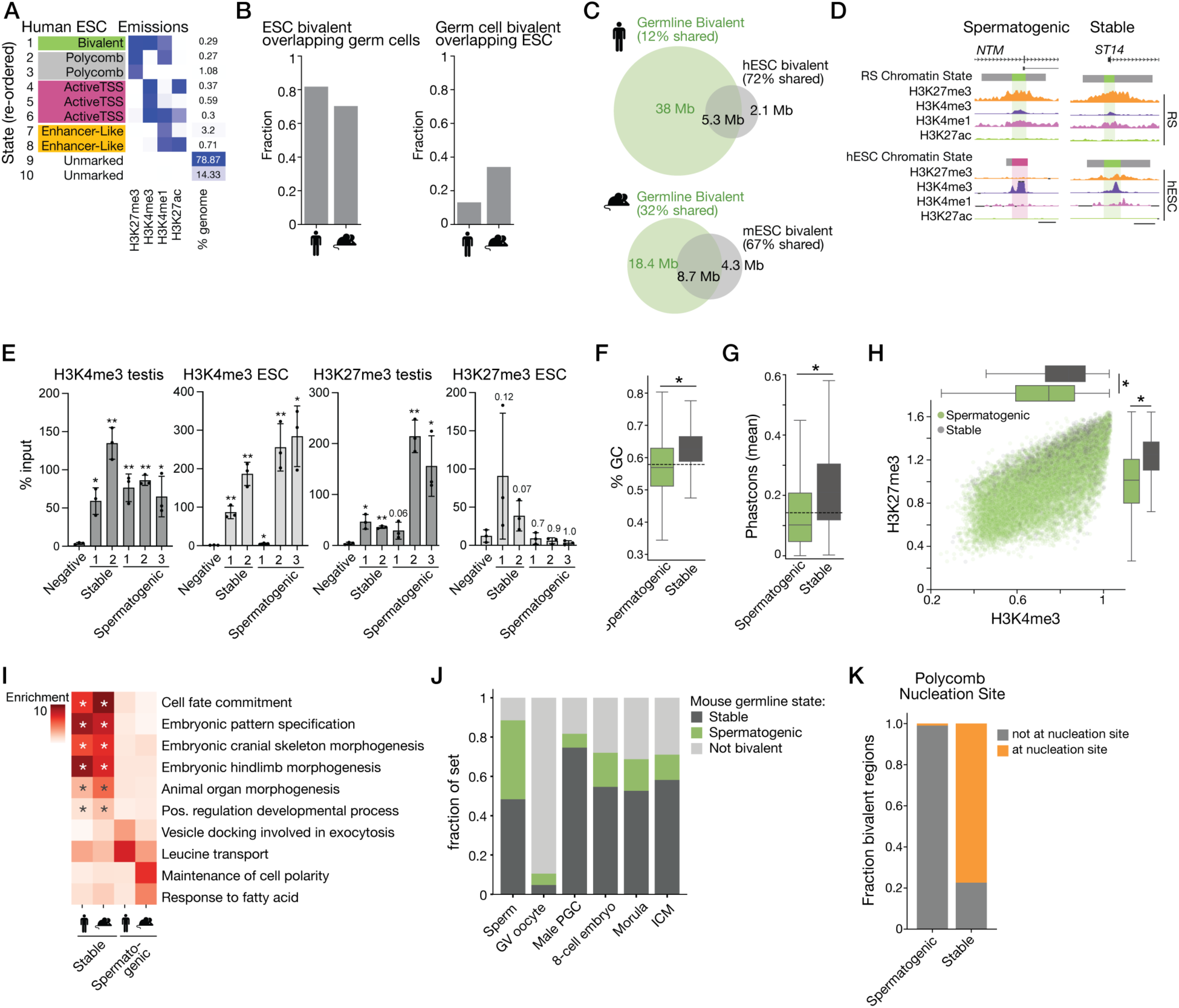
Bivalent chromatin is expansive in male germ cells compared to other in vitro and vivo contexts. **A,** ChromHMM emissions plot and genomic coverage from human ESC ChIP-seq datasets. **B,** Left: fraction of ESC bivalent regions (human, n=6,095, mouse, n=9,900) also bivalent in male germ cells. Right: Fraction of male germ cell bivalent regions (human, n=28,478, mouse, n=18,327) also bivalent in ESC. **C,** Venn diagram representing total genomic coverage (in megabases) of the bivalent chromatin state in human (top) and mouse (bottom) germ cells and ESCs. **D,** Example genome browser tracks of spermatogenic and stable bivalency classes. For spermatogenic bivalent regions (left), the corresponding genomic region is not bivalent in ESC. For stable bivalent regions, the corresponding genomic region is bivalent in ESC. Scale bar equals 2 kb. **E,** H3K4me3 (left) and H3K27me3 (right) ChIP-qPCR in mouse whole testis and mESC samples at two stable bivalent and three spermatogenic bivalent loci.

To understand why some genomic regions acquire bivalency only in germ cells, we subdivided our germline bivalent regions according to their bivalency status in ESCs (**Fig. 3D**, **S3C**), defined as stable regions (bivalent in both spermatogenic and ESCs) and spermatogenic regions (bivalent only in spermatogenic cells) (**Table S3**). We confirmed bivalency at representative stable and spermatogenic bivalent loci using ChIP-qPCR in ESCs and testis tissue (**Fig. 3E**) and compared genomic features of these two groups of bivalent elements. Overall, stable bivalent regions had “stronger” features typical of bivalency compared to spermatogenic bivalent regions, including higher GC content (**Fig. 3F**, **S3D**) and greater sequence conservation (**Fig. 3G**, **S3E**). Stable bivalent elements also had overall higher H3K4me3 and H3K27me3 signal (**Fig. 3H**, **S3F**). In both human and mouse, approximately 80% of stable bivalent regions were promoter-proximal, while spermatogenic bivalent regions were less frequently at promoters (∼50%) (**Fig. S3G**). Genes whose promoters overlap stable bivalent regions were more strongly enriched in developmental gene ontologies (GO), including “cell fate commitment”, “embryonic pattern specification”, and terms related to embryonic morphogenesis (**Fig. 3I**, **Table S4**, Methods). For spermatogenic bivalent regions, there were fewer statistically significant GO terms in either human or mouse, and enriched GO terms were generally not shared across species. Notably, both stable and spermatogenic bivalent regions were enriched for PRC1, including RING1B (**Fig. S3H**) and H2AK119ub^18,44^ (**Fig. S3I**), a common feature of bivalency^13^.

We next asked if these spermatogenic bivalent regions were present only during late spermatogenesis, or if these loci were also bivalent in other germline cell types, including mature sperm, oocytes, male primordial germ cells (PGCs), and preimplantation embryos. We defined bivalent promoters from public datasets in these other cell types in mouse ^5,6,17,45^ and compared them to our stable and spermatogenic bivalent intervals. We found that nearly all bivalent promoters in sperm (89%) are also bivalent in our mouse post-meiotic germ cell maps, with ∼40% of these corresponding to our spermatogenic bivalent set (**Fig. 3J**). In contrast, there was little overlap between oocyte bivalent promoters and bivalent loci in our post-meiotic male germ cells. The majority of bivalent regions in male PGCs and in preimplantation embryos were also bivalent in mouse spermatogenic cells, but most of these were categorized as stable rather than spermatogenic bivalent regions (75% PGC, 55% 8-cell embryo, 53% morula, 58% ICM), reflecting a more ESC-like overall bivalency state. These results indicate that spermatogenic bivalent regions defined in our analysis are distinct from typical bivalent regions defined in ESC studies and their bivalent state is limited to spermatogenic cells. In contrast, stable bivalent regions represent more traditionally defined bivalent loci and appear to retain bivalency throughout germline development.

Previous work identified a set of bivalent domains (“Class II”) that is established de novo in meiotic spermatocytes and distinct from bivalent domains that are present in early spermatogonia and maintained throughout spermatogenesis^19^. Class II bivalent domains are not enriched at developmental promoters, reminiscent of our spermatogenic bivalent gene set. We found striking and specific enrichment for our spermatogenic bivalent regions among the Class II bivalent domains (**Fig. S3K**), where 67% of Class II bivalent genes (501 genes) were classified as spermatogenic bivalent in our data, compared to only 15% of early-established (Class I) bivalent genes (321 genes).

We conclude that a large class of atypical bivalent domains is established during late spermatogenesis and maintained in mature sperm, but largely lost after fertilization.

Bars show mean percent input; error bars show standard deviation between n=3 biological replicates. *p < 0.05, **p < 0.01, 1-sample t-test for each target % input compared to mean % input at the negative locus (alternative=”greater”). **F,** GC content in bivalent regions grouped by ESC status. Box shows median and interquartile range; whiskers show range of the distribution. Dashed line indicates mean GC content for of all human bivalent regions. *p < 1e-30, two-sample, two-sided Kolmogorov-Smirnov (KS) test. **G,** Mean phastCons score within individual regions by ESC status. Dashed line indicates the phastCons mean value for of all human bivalent regions. *p < 1e-30, two-sample, two-sided KS test. **H,** Quantitative H3K4me3 and H3K27me3 ChIP-seq signal in human bivalent regions color-coded by hESC status (spermatogenic, n=24,593; stable n=3,719). *p < 1e-30, two-sample, two-sided KS test. **I,** Selected gene ontology terms and their enrichment level within the sets of gene promoters associated with stable bivalency (left) or spermatogenic bivalency (right). Genes were associated with bivalent domains if the domain overlapped an interval +/-1kb from the TSS. (Number of genes: human stable n=2,003; mouse stable n=3,200. Human spermatogenic n=4,809; mouse spermatogenic n=2,724). *Adjusted p-value < 0.05. **J,** Fractions of bivalent promoters defined in other cellular contexts with a given bivalency status in the mouse germline. Total bivalent promoters considered: sperm^45^ n=3,085, GV oocyte^17^ n=6,095, male PGC^5^ n=694, 8-cell embryo n=1,933, morula n=1,211, ICM n=2,105. Embryo bivalency data obtained from Liu et al^6^. **K,** Fractions of mouse stable or spermatogenic bivalent regions overlapping Polycomb nucleation sites (Veronezi et al^46^, Methods). Total bivalent regions considered: spermatogenic n=12,077, stable n=6,250.

### Spermatogenesis-specific bivalent regions do not require strong Polycomb nucleation sites

The finding that spermatogenic bivalent regions have weaker bivalent features implies that the chromatin environment of late spermatogenic cells creates a more permissive threshold for establishment of bivalency at a given sequence compared to ESCs and other germline cell types. We confirmed that the core PRC2 and PRC1 machinery is present in both ESCs and spermatogenic cells, indicating that presence or absence of the appropriate histone modifying activity does not explain this difference in threshold (**Fig. S3L**). As a further test, we also compared our bivalent region sets to Polycomb nucleation sites defined in ESCs ^46^. We found that whereas 77% of stable bivalent regions overlapped nucleation sites, only 1% of spermatogenic bivalent regions did (**Fig. 3K**). This finding implies that the late spermatogenic chromatin environment establishes and regulates bivalency differently from ESCs and other germline cells.

### Bivalent regions in spermatogenic cells are associated with distinct evolutionary trajectories

To elucidate the unique bivalency-regulating features of spermatogenic cells, we leveraged our cross-species epigenomic datasets to identify sets of bivalent regions with different chromatin evolutionary histories, predicting that regions with evolutionarily recent gains of bivalency in late spermatogenic cells share features that promote spermatogenic bivalency. We classified regions based on sequence orthology between species and compared ChromHMM-defined chromatin states at orthologous regions (Methods). To minimize the problem of noise affecting a specific species, we focused our analysis on regions where the chromatin state was conserved within the primate (both human and rhesus) or rodent (both mouse and rat) lineages, and excluded ancestral elements that had discordant chromatin states in bull and opossum (Methods). For clarity, we use the term “ancestral” to refer to the chromatin state in bull and opossum, inferred to approximate the ancestral state, although we did not use phylogenetic signal to infer ancestry. Using these annotations, we quantified the frequency of different chromatin evolutionary trajectories across species in germ cells (**Fig. 4A**, **S4A**). As expected, the largest sets of regions showed conservation of active chromatin (35% of classified regions in primate, 25% in rodent), followed by conserved Polycomb chromatin state (23% of classified regions in primate and 21% in rodent). Bivalency was conserved at a smaller but substantial fraction of regions (5.5% of classified regions in primate, 8.6% in rodent), consistent with its lower overall prevalence in the genome. Regions where bivalency was gained in either rodent or primate lineages also represented a relatively large class, while we rarely captured bivalency losses. We focused further analysis on regions where bivalency status was either conserved (‘conserved bivalent’) or gained (‘ancestrally Polycomb’ and ‘ancestrally active’) (**Fig. 4B-C**, **S4B**, **Table S5**).

**Figure 4:**
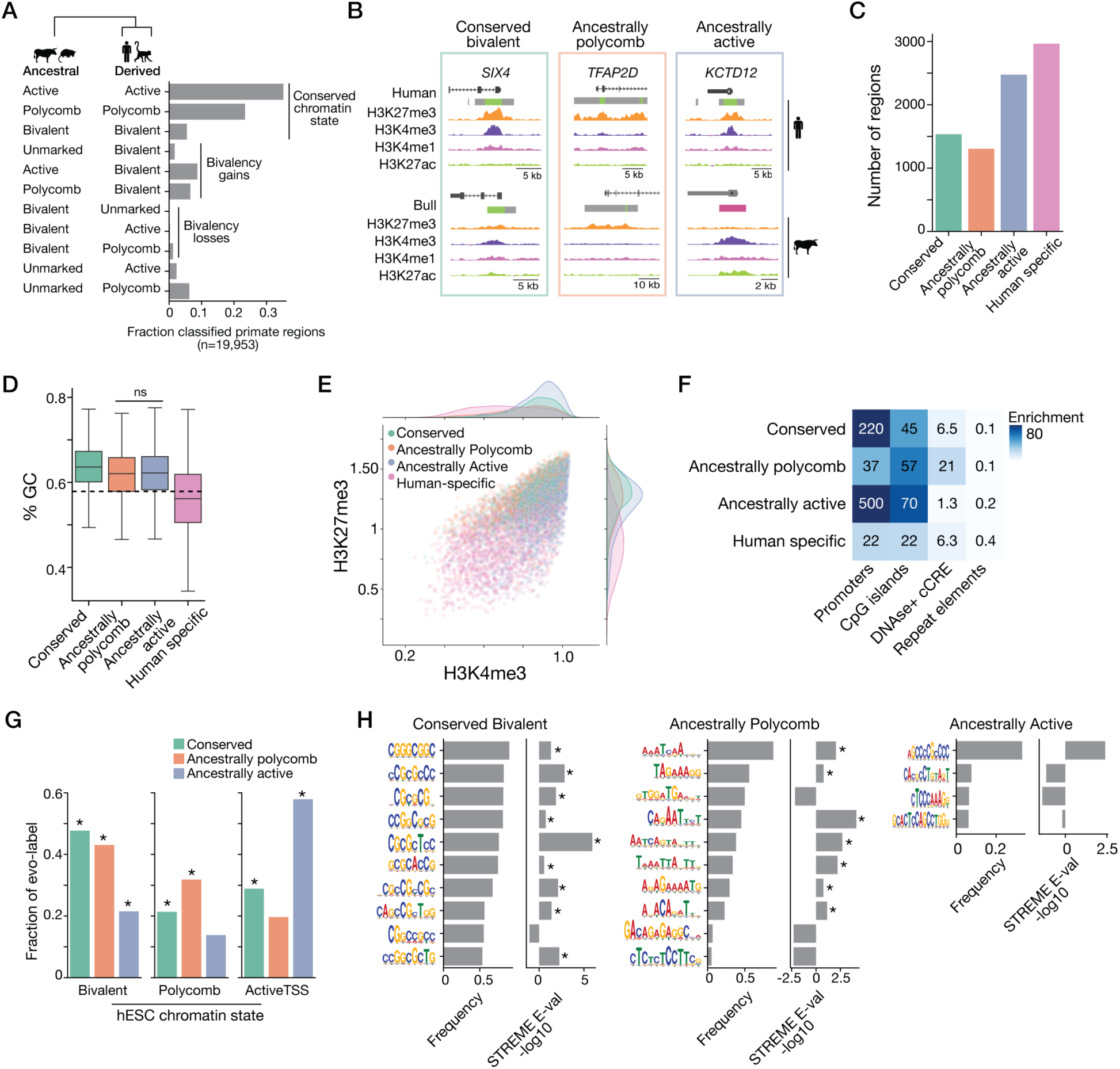
Male germ cell bivalent regions have homogenous chromatin features but enrichment for distinct DNA sequences based on ancestral chromatin state. **A,** Frequencies of orthologous chromatin state combinations between ancestral species (bull, opossum; left) and primate species (rhesus, human; right). The most frequent states are shown based on n=19,953 total regions classified. **B,** Example genome browser tracks of representative examples of evolutionarily dynamic bivalency classes: conserved bivalent (left), ancestrally Polycomb (middle), and ancestrally active (right). **C,** Numbers of bivalent regions in each set of evolutionary classes defined in the primate lineage. **D,** GC content of individual human regions within each class of evolutionarily dynamic bivalent regions. Box shows median and interquartile range; whiskers show range of the distribution. Dashed line indicates mean GC content for all bivalent regions. All comparisons significantly different (p < 1e-30, two-sample, two-sided Kolmogorov-Smirnov test) except where indicated as “ns”. **E,** Quantitative H3K4me3 and H3K27me3 ChIP-seq signal in human bivalent regions color-coded by evolutionary class. Number of elements in each class as shown in (C). **F,** Genomic enrichment (odds ratio) of promoters, CpG islands, DNase+ H3K4me3 cCRE elements, and repeat elements (LINE/SINE/LTR) in distinct evolutionary classes of human bivalent elements. **G,** Fraction of human bivalent regions within an evolutionary class that have the indicated chromatin state in hESCs. Total number of elements within each set as shown in (C). *p < 0.05, Fisher’s exact test (alternative=’greater’) for comparison of co-occurrence between states. **H,** Position weight matrix of the most frequent enriched DNA-motifs found in human bivalent regions according to evolutionary label, with total number of elements as shown in (C), using all evolutionary classified regions as negative background set. For each enriched motif, frequency in the test set (left) and motif E-value (-log10) (right) is shown (Methods). *p < 0.05.

We first asked if regions which are all considered bivalent but have different evolutionary histories exhibited distinct genomic and functional features. Overall, we found that conserved, ancestrally active, and ancestrally Polycomb bivalent regions had similar genomic features, including GC content (**Fig. 4D**, **S4C**), H3K4me3 and H3K27me3 enrichment levels (**Fig. 4E**, **S4D**) and genomic enrichment at regulatory regions (**Fig. 4F**, **S4E**). For comparison, we also defined very recently gained ‘species-specific’ bivalent regions (e.g. bivalent only in human and not rhesus) and found that these had notably lower GC content and histone modification enrichment signal. Species-specific and ancestrally Polycomb bivalent regions were also less enriched at promoters, and ancestrally Polycomb bivalent regions were more frequently observed at open chromatin regions elements (ENCODE defined DNase sensitive regions) compared to conserved and ancestrally active bivalent regions (**Fig. 4F**, **S4E**). We observed very low frequencies of overlap with repeat elements, with no meaningful differences between bivalent regions classified by ancestral chromatin status. We conclude that bivalent regions with deeply conserved or lineage-specific bivalency exhibit similar overall sequence and genomic features, while very recent species-specific bivalent regions are more weakly associated with these features, possibly representing emergent regions of bivalency.

Given the overall similar genomic features between evolutionary classes, we next asked if conserved, ancestrally Polycomb, and ancestrally active bivalent regions also retained bivalency to a similar extent in ESCs. Many bivalent elements in all three classes are also bivalent in ESCs (**Fig. 4G**, **S4F**). However, we observed a striking correspondence between ancestral germline chromatin state and ESC chromatin state. Specifically, conserved bivalent regions most frequently exhibit stable bivalency in ESCs, ancestrally active bivalent regions are more often marked with active chromatin state ESCs (58%), and ancestrally Polycomb bivalent regions are more often marked with the Polycomb chromatin state (32%).

### Differential motif enrichment in bivalent domains corresponds to chromatin evolutionary history

Although overall genomic features of conserved, ancestrally active, and ancestrally Polycomb bivalent domains were globally similar, we hypothesized that bivalent domains arising from different ancestral chromatin states might follow distinct molecular pathways to gaining bivalency that leave detectable signatures revealed in class-specific sequence motifs. To test this, we performed another de-novo DNA sequence motif enrichment analysis for each evolutionary class of bivalent regions. We used bivalent sequence from all three evolutionary labels as the background to identify motifs unique between sets and avoid general bivalency-enriched motifs (Methods). Surprisingly, despite their homogenous chromatin features, we observed consistent differences in enriched motif sets between bivalent regions with distinct evolutionary histories (**Fig. 4H**, **S4G**). In conserved bivalent elements, enriched motifs were dominated by 8-10 base-pair-long CG-rich motifs, similar to bivalent elements as a whole (**Fig. 1K**). In contrast, sequences enriched specifically in ancestrally Polycomb bivalent regions were dominated by AT-rich motifs. These results suggest that one mechanism by which Polycomb-dominant regions can gain bivalency over evolutionary time is by acquisition of AT-rich sequence motifs, consistent with in vitro studies where increasing AT sequence composition disrupts bivalency establishment at GC-rich promoters by tipping the balance between GC-dependent Polycomb activity and AT-responsive activating transcription factors ^47^. In ancestrally active bivalent elements, only four motifs were identified, and these were longer (10-15 bp), lower in frequency, and more variable in composition compared to conserved bivalent and ancestrally Polycomb motifs, suggesting that active chromatin elements could gain bivalency through varied molecular mechanisms and associated sequence trajectories. We conclude that although evolutionarily distinct bivalent regions are homogenous in their chromatin and global sequence features, they are distinct at the sequence level based on specific classes of DNA sequence motifs.

### Regulation in somatic tissues recapitulates ancestral germline chromatin state

We next asked if genes associated with different evolutionarily categories of bivalent elements exhibit distinct expression patterns in non-germ cell contexts. We linked genes to nearby (+/-1kb of TSS) bivalent regions and assigned genes to an evolutionary category if they were uniquely associated with regions in a single evolutionary category (Methods, **Fig. S5A**, **Table S6**). We observed a remarkably consistent expression pattern across ESCs (**Fig. 5A**), pre-implantation development (**Fig. 5B**), and adult somatic tissues (**Fig. 5C-D**, **S5B-C**). Genes associated with conserved and ancestrally Polycomb bivalent regions had low expression across all tissues and cell types, while ancestrally active regions had higher expression. This finding suggests that despite the similarity of their chromatin state in spermatogenic cells, these regions have distinct transcriptional regulatory capacities corresponding to their inferred ancestral germline chromatin state. Interestingly, this difference was least apparent in very early embryonic stages (2-cell to 8-cell, **Fig. 5B**), re-emerged in the morula and epiblast, and persisted in all adult somatic tissues examined.

**Figure 5:**
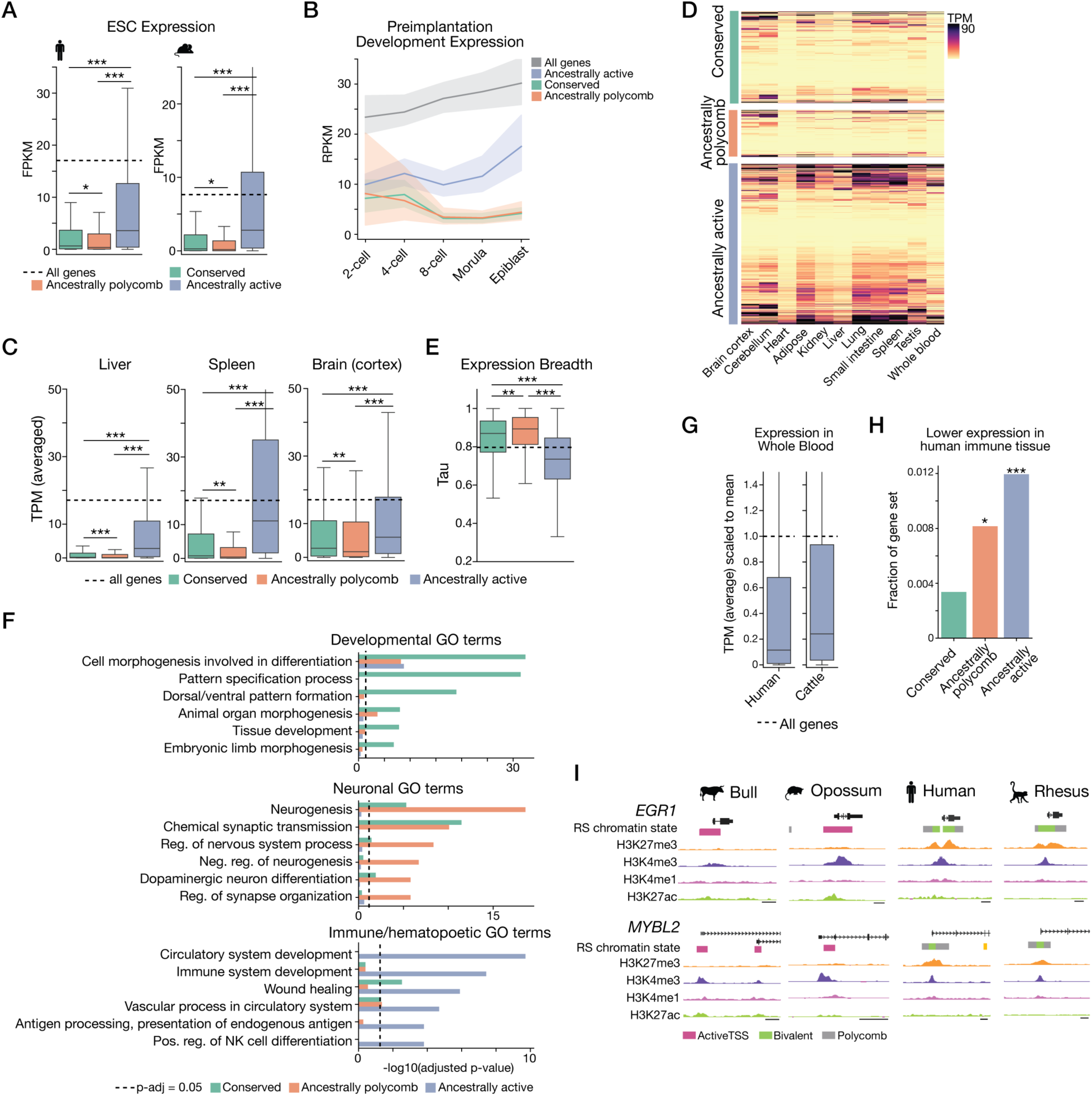
Bivalent regions with different evolutionary histories regulate genes with distinct functions. **A,** Expression level (ENCODE) in human and mouse ESCs for genes assigned to each bivalent evolutionary class. Box shows median and interquartile range; whiskers show range of the distribution. Dashed line shows the mean expression of all genes in ESCs. *p < 0.05, ***p < 0.001, two-sample, two-sided Kolmogorov-Smirnov (KS) test. **B,** Expression levels (RPKM) across human pre-implantation developmental stages^50^ for genes associated with each evolutionary class. Line indicates mean value; shaded area shows 95% confidence interval. Expression data obtained from Yan et al, 2013. **C,** Expression level (mean TPM from GTEx; see Methods) of genes associated with each evolutionary class in select human adult somatic tissues: liver, spleen, and cortex. Dashed line shows mean expression of all genes within the tissue type. **p < 0.01, ***p < 0.001, two-sample, two-sided KS test. **D,** Heatmap of gene expression across adult somatic tissues sampled in GTEx for the three sets of evolutionarily classified genes. Rows are genes and columns are tissues. **E,** tau, a measure of tissue expression breadth, for genes associated with each evolutionary category. Dashed line shows mean tau value of all genes. **p < 0.01, ***p < 0.001, two-sample, two-sided KS test. **F,** Selected developmental (top), neuronal (middle) and immune (bottom) gene ontology (GO) terms and the Benjamini–Hochberg adjusted p-value (-log10) for enrichment of the GO term in each evolutionary gene set category. Dashed line indicates adjusted p-value = 0.05. **G,** Expression level of genes in the ancestrally active evolutionary category (n=2,139) in human whole blood (GTEx, left) compared to bull whole blood (CattleGTEx, right; Methods). Dashed line indicates average expression across all genes in whole blood. Data are normalized to the mean expression level. **H,** Fraction of each gene set by evolutionary category that shows lower expression in human compared to cattle in at least one immune relevant cell type or tissue (CD8+ T cells, CD4+ T cells, spleen, or bone marrow; Methods). *p < 0.05, ***p < 0.001, Fisher’s exact test, alternative=’greater’, indicating the likelihood of the observed number of genes occurring in each set by chance. (conserved n=1202, ancestrally Polycomb n=624, ancestrally active n=2139). Differentially expressed genes (n=256) obtained from Yang et al, 2022^51^. **I,** Genome browser showing germline chromatin state of *EGR1* and *MYBL2* in bull, opossum, human, and rhesus. Scale bar equal to 2 kilobases.

Since different germline ancestral states predicted distinct somatic expression profiles, we next asked if these gene sets also differed in other characteristics. We found that genes associated with conserved bivalent and ancestrally Polycomb states are more tissue specific as assayed by a metric for tissue expression breadth (tau^48,49^) (Methods), while genes associated with ancestrally active bivalent regions were more broadly expressed (**Fig. 5D-E**, **S5C-D**).

We next performed a gene ontology (GO) enrichment analysis for these gene sets in human (**Fig. 5F**) and mouse (**Fig. S5E**, Methods), and found that evolutionarily distinct gene sets were enriched for functions in distinct somatic tissues. Genes associated with conserved bivalent domains were enriched for general developmental GO terms, including “pattern specification processes,” “cell fate commitment” and “animal organ morphogenesis”, consistent with typical regulatory targets of bivalent domains. In contrast, genes associated with ancestrally Polycomb regions showed greater enrichment for neuronal terms, including “neurogenesis” and “regulation of nervous system process”, although some neuronal terms were also significantly enriched in the conserved bivalent gene set. Finally, genes associated with ancestrally active regions showed strong and selective enrichment for immune and hematopoiesis-related GO terms, including “circulatory system development,” “immune system development,” and “positive regulation of natural killer cell differentiation.” These patterns were consistent in human and mouse data, although neuronal terms were slightly less specific to ancestrally Polycomb genes in mouse compared to human. Together, we find that evolutionarily distinct classes of bivalent regions regulate functionally different gene sets and exhibit different expression patterns across somatic tissues, despite having indistinguishable, homogenous regulatory states in the male germ line. This suggests that evolutionary gains of bivalency in the germline demarcate regulatory changes that preferentially impact certain somatic tissues in primate and rodent species depending on their initial ancestral chromatin state.

### Divergent expression and function in immune tissues correlates with gain in spermatogenic bivalency

Since we observed that ancestrally active bivalent genes in both the primate and rodent lineages were strongly enriched for immune-relevant gene functions, we hypothesized that germline gains in bivalency in primates and rodents may mark genes whose regulatory state and function in immune cells underwent recent evolutionary change in these lineages.

Specifically, transition from an active to a bivalent chromatin state in germ cells may correspond to sequence changes that reduce overall transcriptional activity, tipping the balance toward increased H3K27me3 in germ cells as well as decreased expression in immune cells. To test this hypothesis, we evaluated the expression of the ancestrally active gene set in human and cow whole-blood samples, obtained from GTEx and cattleGTEx^52^. Since quantitative comparisons in these independent datasets from different species are not robust on a gene-by-gene basis, we used expression of all genes normalized to the mean expression level in blood as a comparative guide. We observed that the expression of ancestrally active genes in human blood was overall lower than in cattle (**Fig. 5G**). This finding is consistent with a model where germline gains of bivalency mark genes whose transcriptional activity is reduced in the relevant somatic tissues. Next, we examined a set of genes reported to be differentially expressed between cattle and human immune tissues^51^ and asked if bivalent genes classified as ancestrally active in human spermatogenic cells were overrepresented among genes with lower expression in human compared to bovine immune cells. Indeed, of the 256 genes with lower expression in human, twenty-five were ancestrally active bivalent genes (p < 0.001, Fisher’s exact test) (**Fig. 5H**). Among these were *MYBL2* and *EGR1*, each of which has key roles in immune development or function (**Fig. 5I**). *EGR1* is important for human myeloid cell differentiation^53^ and inflammatory response^54^, and loss of *MYBL2* leads to depletion of the hematopoietic stem cell pool^55^. In contrast, the fraction of ancestrally active genes was lower among genes with increased or conserved expression levels between cattle and human, although still statistically enriched, reinforcing the relationship between immune function and ancestrally active promoter chromatin states (**Fig. S5G**).

## Discussion

Since its initial discovery, the bivalent chromatin state has been an intriguing yet functionally enigmatic regulatory feature of cells with pluripotent potential. Here, we comprehensively characterize bivalency in mammalian male germ cells to reveal a potential role for bivalency in balancing cooccurring demands on regulatory sequence, where the same sequence must both support germ cell function and direct somatic gene regulation. By utilizing a comparative cross-species and developmental epigenetic framework, we identify a unique bivalent chromatin landscape in spermatogenic cells. We find that spermatogenic bivalent chromatin has features similar to ESC bivalency, but is also present at loci less typically associated with bivalency, including intergenic regions and promoters of non-developmental genes. We further identify evolutionarily dynamic loci where bivalency has been gained, and find that genes with evolutionary gains of bivalency in the primate and/or rodent lineages have distinct expression profiles in somatic tissues even though they have largely indistinguishable bivalent regulatory states in the germline. Our results suggest that gain of bivalency from ancestrally active sequence elements is selectively associated with functional changes in immune cells. We propose that bivalency in male germ cells functions in part to allow regulatory sequence to evolve while protecting spermatogenic cell function (**Fig. 6**).

**Figure 6.**
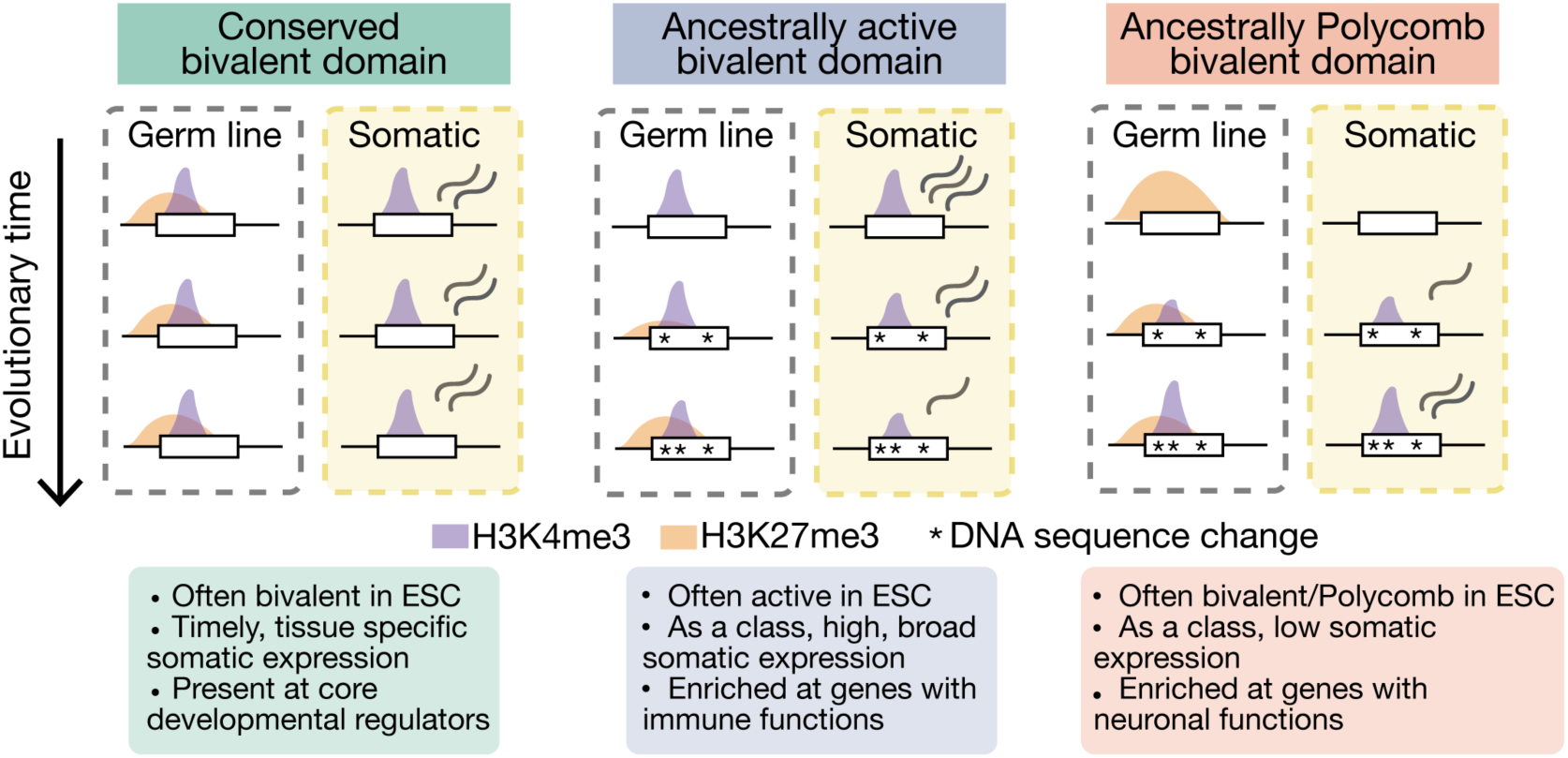
Model for evolution of bivalent domains in the germline and associated regulatory impact in somatic tissues.

A striking finding in our analysis is the markedly elevated amount of bivalent chromatin in spermatogenic cells compared to ESCs (**Fig. 3C**). This suggests that (1) spermatogenic cells are more permissive to bivalency establishment or (2) there may be unique, spermatogenic-specific factors controlling deposition of bivalent chromatin. Our analysis suggests that less favorable genomic sites can establish Polycomb in spermatogenic cells (**Fig. 3K**), although direct measurement of PRC2 nucleation in spermatogenic cells would be informative. We verified that the same chromatin machinery canonically known to direct bivalency is present in both testis and ESCs (**Fig. S3L**), although we have not excluded the possibility of differential activity or fine-tuned dynamic regulation of these trans-acting factors. Previous work identified SCML2 as a germline-specific PRC1 cofactor important for bivalency establishment^19^. Using public data, we confirmed that SCML2 is present in pachytene cells at both spermatogenic and stable bivalent loci (**Fig. S3J**), suggesting that this factor is generally supportive to bivalency establishment in spermatogenic cells but does not distinguish stable and spermatogenesis-specific bivalent sites. Future work is needed to understand the molecular mechanisms supporting establishment of spermatogenic bivalent chromatin.

We observe that most bivalent promoters in epididymal sperm^45^ are also bivalent in our post-meiotic datasets. In contrast, many spermatogenic bivalent regions do not retain bivalency in other developmental contexts, including the preimplantation embryo and ICM (**Fig. 3J**). This observation suggests that spermatogenic bivalent regions likely do not function to transmit continuous regulatory information into the embryo, but instead play an important role in the biology of spermatogenic cells themselves. This phenomenon of coexisting “stable” (more typical, present in ESCs) and less stable or atypical bivalent regions has been reported in other cell types, including the epiblast and the peri-implantation embryo^56,57^. This pattern may suggest a general feature of bivalent chromatin regulation, where bivalency consistently targets a set of typical genomic loci, as well as a separate, variable set of atypical regions that are uniquely bivalent in a given cellular context and may play context-specific roles. In spermatogenic cells, broad bivalency establishment could be a response to the high transcriptional complexity of the testis, which is particularly relevant in meiotic and post-meiotic cell types^28^. Transcriptional promiscuity in the testis has been cited to suggest that spermatogenic cells are an evolutionary “testing ground”, and in this context bivalent chromatin could act to minimize or offset deleterious impacts of new transcripts or regulatory features that impair spermatogenic function and viability but have an advantageous effect in the soma. In support of this model, loss of some germline bivalency regulators, including SCML2 and MLL2, results in disruption of spermatogenesis, indicating that maintenance of bivalency is important for germ cell function and viability ^44,58^. However, as both SCML2 and MLL2 regulate stable bivalent regions, this phenotype also does not separate the functions of stable and spermatogenic bivalent regions.

We found strong enrichment for immune-related functional categories among genes that gained bivalency in the primate or rodent lineages from an inferred ancestrally active state (**Fig. 4F, S4E**). This finding suggests that the germline regulation of genes important in the immune system, more than other somatic tissues, may change rapidly across species. Immune genes often exhibit signs of positive selection in their coding sequence ^59^; those involved in pathogen recognition, such as cell surface proteins and cytokine receptors, have particularly high positive selection signals ^60^. In tandem with coding sequence, regulation of these genes may also be evolving rapidly. However, measuring signatures of evolutionary selection on non-coding functional regions is difficult^61^. We propose that regulatory evolution occurs rapidly at immune genes, analogous to and sometimes in coordination with coding changes, and these changes can create targets for spermatogenic-specific bivalency establishment. We recently reported this phenomenon at the *Traf6* promoter, where DNA sequence divergence in the rodent lineage resulted in both differential regulation of this gene in the immune context and gain of the bivalent state in the male germline^62^. More generally, we propose that evolutionary gains of bivalent chromatin in the germline demarcate elements undergoing regulatory change, where the impact of these changes will be fully realized in somatic tissues. Spermatogenic cells may thus utilize bivalency as a method to protect germ cell function from the impacts of newly arising regulatory changes that confer an advantage to the organism as a whole.

## Methods

### mESC culture

Mouse embryonic stem cells were grown on 0.1.% gelatin (EmbryoMax, Millipore ES-006-B) and mouse embryonic fibroblast (MEF) feeder cells (Gibco MitC-treated CF-1 MEF A34959) at 37°C and 5% CO2 in a humidified incubator in embryonic stem cell media (DMEM (Gibco 11965-092) with 15% fetal bovine serum (Sigma F4135), Penicillin-Streptomycin (1:100; Gibco 15140-122), GlutaMax (1:100; Gibco 35050-061), MEM NEAA (1:100; Gibco 11140-050), Sodium Pyruvate (1:100; Gibco 11360-070), HEPES (1:40; Gibco 15630-080), β-mercaptoethanol (1:125,000; Sigma-Aldrich M6250), and LIF (1:10,000; Millipore ESG1106)). Media was changed daily and cells were passaged every 2-3 days with StemPro Accutase (Gibco A11105-01). Growing cells were tested for mycoplasma every 1-2 months to ensure lack of contamination. For Western blot, ESC lysates were prepared using RIPA lysis buffer (G Biosciences 786-489) and supplemented with protease inhibitor cocktail (Sigma Aldrich 11836170001).

### Germ cell collection for mouse and rat ChIP

Whole testes were dissected out of the animal and placed into a petri dish with PBS. Testes were decapsulated and dissociated as previously described^63^. Briefly, decapsulated testis were added to 0.75 mg/mL Type IV collagenase (Gibco 17104-019) in DMEM and incubated at 37°C for 10 minutes, with gentle inversion and pipetting to dissociate into spaghetti-like strands. The collagenase reaction was quenched by adding an equal volume of DMEM and cells were pelleted at 400xg for 5 minutes at room temperature. Cells were washed with DMEM, then resuspended in 0.05% trypsin (Gibco 25-200-056) and incubated for 5-10 minutes. The trypsin reaction was quenched by adding an equal volume of DMEM + 10% Calf Serum (Sigma C8056). Cells were pelleted as before and washed twice with DMEM + 10% CS. The suspension was filtered through a 100 uM filter. Round spermatids were isolated by flow cytometry as previously described^63^. Briefly, dissociated testis cells were incubated at room temperature with 2 µl/ml Vybrant DyeCycle Green (Invitrogen V35004) for 30 minutes. Round spermatids were sorted based on 1C DNA content and size using a 2-laser sorter (Bio-Rad S3e with 488nm and 561nm lasers) gated on single cells.

### Testis lysates for Western blot

Testes were dissected from mice and the tunica albuginea was removed. A single cell suspension was formed by digesting the testis with collagenase IV (Gibco17104-019) and then with 0.05% trypsin (Gibco 25300-054) at 37°C for 10 minutes each. Lysates were then prepared from this single cell suspension using RIPA lysis buffer (G Biosciences 786-489) and supplemented with protease inhibitor cocktail (Sigma Aldrich 11836170001).

### Chromatin immunoprecipitation

Chromatin immunoprecipitation was performed as previously described^64^. 1-3e6 cells were fixed in methanol-free formaldehyde to a final concentration of 1%, with rotation at room temperature for 10 minutes. Cold 1.25 M glycine was added to quench the reaction to a final concentration of 125 mM. Cells were pelleted at 1,500xg for 5 minutes at 4°C. The pellet was washed with cold PBS twice, followed by centrifugation as before. Cells were resuspended in 300 uL ChIP lysis buffer (800 mM NaCl, 25 mM Tris pH 7.5, 5 mM EDTA pH 8, 1% v/v Triton X-100, 0.1% w/v SDS, 0.5% w/v sodium deoxycholate), snap frozen in liquid nitrogen, and stored at −80C.

For each ChIP, 20 uL of Protein G DynaBeads (Invitrogen 10003D) were resuspended in 40 uL chromatin dilution buffer (25 mM Tris pH 7.5, 5 mM EDTA pH 8, 1% w/v SDS) and incubated with rotation at room temperature with 1-10 uL of antibody for 3 hours (antibodies used are listed in **Table S7**). Samples were thawed on ice and sonicated at 4°C for 10 cycles (30 seconds on/off) using a Bioruptor (Diagenode). 1 mL of protease inhibitor supplemented chromatin dilution buffer (1X, Roche 11873580001) was added to the sonicated samples, which were then centrifuged at 4°C for 30 minutes at 13,600xg. The soluble chromatin was moved to a fresh tube, and 1-5% of material was set aside at −20°C to use as input. Dynabeads coupled to the antibody were added to the soluble chromatin and incubated with rotation overnight at 4°C. The next day, beads were collected using a magnet and unbound chromatin was discarded.

Beads were washed once with Wash Buffer A (140 mM NaCl, 50 mM HEPES ph 7.9, 1 mM EDTA pH 8, 1% v/v Triton X-100, 0.1% w/v SDS, 0.1% w/v sodium deoxycholate, adjusted to pH 7.9), Wash buffer B (500 mM NaCl, 50 mM HEPES ph 7.9, 1 mM EDTA pH 8, 1% v/v Triton X-100, 0.1% w/v SDS, 0.1% w/v sodium deoxycholate, adjusted to pH 7.9), Wash buffer C (20 mM Tris pH 7.5, 1 mM EDTA pH 8, 250 mM LiCl, 0.5% w/v NP-40 Alternative, 0.5% w/v sodium deoxycholate, adjusted to pH 8), and two times with TE (10 mM Tris pH7.5, 1 mM EDTA pH 8, 1% w/v SDS). Elution from the beads was performed twice by adding 100 uL fresh elution buffer (10 mM Tris pH 7.5, 1 mM EDTA pH 8, 1% w/v SDS) to the beads, followed by 5 minute incubation at 65°C, followed by 15 minutes incubation at 65°C with agitation. Eluted DNA was moved to a fresh tube and NaCl was added to samples to a concentration of 160 mM. Samples were treated with RNase A (20 uG/mL, Millipore Sigma) and incubated at 65°C for at least 8 hours to remove RNA contamination and reverse crosslinking. After RNase A treatment, EDTA was added to samples to a concentration of 5 mM. Proteinase K (NEB) was added to a final concentration of 200 uG/mL and incubated at 45°C for 2 hours. ChIP DNA was purified using the Zymo ChIP DNA Clean & Concentrator kit (Zymo D5201) following manufacturer’s instructions. Purified ChIP-DNA was used for sequencing library preparation or for ChIP-qPCR. For ChIP-seq, Illumina 150 bp paired-end sequencing libraries were generated.

### qPCR

Quantitative PCR was performed in 20 uL reaction volume (1 uL ChIP DNA, 10 uL Power SYBR Green PCR Master Mix (Applied Biosystems, 4367659, Thermo Fisher Scientific, Inc.), 8 uL nuclease-free water, and 1 uL of 10 uM forward and reverse primer mixture). Reactions were performed in technical triplicate in a 96 well plate using a QuantStudio 3 Real-Time PCR System (Applied Biosystems, Thermo Fisher Scientific, Inc.) with standard cycling conditions (Hold stage (x1): 50°C for 2 minutes, 95°C for 10 minutes; PCR stage (x40): 95°C for 15 seconds, 60 °C for 1 minute, Melt curve stage: 95°C for 15 seconds, 60°C for 1 minute, 95°C for 15 seconds). Enrichment at a specific locus was calculated as a percentage of total input chromatin using the following equations:

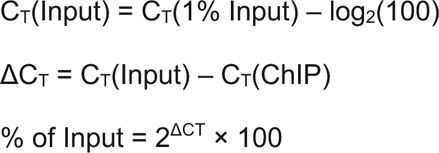

Primer sequences are provided in **Table S7.**

### Western Blot

10 micrograms of whole testis or ESC lysate was diluted with 4X Laemmli Buffer (BioRad 1610747), boiled at 95°C for 5 minutes, and then loaded on to a 4-20% gradient stain-free protein gel (BioRad 4568093). After gel electrophoresis, the proteins were transferred to a 0.45 uM nitrocellulose membrane (BioRad 1620115) and blocked for 1 hour at room temperature with 5% milk (RPI M17200) in TBST (RPI T59180). Membranes were then incubated in primary antibody overnight at 4°C (specific antibodies and dilutions provided in **Table S7**). The next day, an anti-rabbit-HRP conjugated secondary (Vector Laboratories, P1-1000) was used at 1:50,000 for 1 hour at room temperature. Signal was detected using SuperSignal™ West Pico PLUS Chemiluminescent Substrate (Thermo Scientific 34580) and a ProteinSimple FluorChem E FE0685 Imager.

#### Analysis methods

Unless otherwise specified, computational programs were used with default parameters.

### ChIP-seq data processing

Sequencing reads were filtered for quality using FASTX-Toolkit fastq_quality_filter (v.0.0.14, -p 80 -q 20), followed by trimming adaptor sequences using CutAdapt^65^ (v3.4). Paired-end reads were paired using SeqKit^66^ pair (v2.3.1). Filtered and trimmed reads were aligned to the relevant genome assembly (hg38, rheMac10, mm39, rn6, mondom5, bosTau9) using Bowtie2^67^ (v2.4.2, --sensitive). Duplicate reads were removed from .bam files using Picard^68^ MarkDuplicates (v2.25.6, --REMOVE_DUPLICATES true). Biological replicates were compared using deepTools^69^ multiBamSummary bins (--region chr3) and correlations were visualized using plotCorrelation (--corMethod pearson --skipZeros --whatToPlot heatmap) (**Fig. S1**).

### Visualization of ChIP-seq data

Bigwig signal tracks were generated from aligned .bam files using deepTools bamCoverage, normalizing for read depth (v3.5.1, --normalizeUsing CPM), and visualized using the UCSC Genome Browser^70^ (smoothingWindow 5). Metagene plots were generated using deepTools computeMatrix (--referencepoint --binSize 100) and plotProfile. For sets of shuffled regions, a random sample of 5,000 regions from the regions of interest were selected and randomly shuffled throughout the genome using BEDTools^71^ shuffle.

### Generating chromatin state maps using ChromHMM

All datasets used in the ChromHMM segmentation are provided in **Table S2**. For each chromatin state map (germ cell chromatin state in human, rhesus, mouse, rat, bull, opossum; embryonic stem cell state in human, mouse), a tab delimited file was created specifying the input datasets, the associated ChIP-seq modification, and cell type of origin. ChIP-seq signal was first binarized using ChromHMM^35,36^ (v1.24) BinarizeBam (binsize = 200bp), followed by ChromHMM LearnModel, using a range of state numbers (8, 10, 12, 14). For human, rhesus, rat, bull, and opossum, the round spermatid segmentation was used. For mouse, germ cell types were not distinguished in the ChromHMM model, and the singular germ cell segmentation was used. To determine optimal state segmentation, states corresponding to bivalent (H3K4me3+, H3K27me3+), activeTSS (H3K27me3-), and Polycomb (H3K27me3+) were defined using emission plots, and bivalent, activeTSS, and Polycomb regions called in each segmentation were visually compared to ChIP-seq signal on the genome browser with the segmentation parameters blinded. The state number which best captured ChIP-seq signal data was chosen for each species and cell type. We found that this corresponded to the 12-state model in all germ cell segmentations and the 10-state model for ESC segmentation.

ChromHMM OverlapEnrichment was used to determine enrichment of individual chromatin states in RefSeq^72^ genome annotations (obtained from the UCSC Table Browser^73^, Track: NCBI Refseq, Table: RefSeq Curated), in bivalent gene promoters as previously reported^14^, and in gene promoters with high and low expression in round spermatids^14^. For mouse and human, enrichment was also calculated in ENCODE defined cCREs^74^ (Registry V2).

For each ChromHMM state, histone modification and genome annotation enrichments were used to assign functional labels: bivalent, Polycomb, activeTSS, enhancer-like, or unmarked. States where manual inspection in the browser indicated that signal was excessively noisy were not assigned a functional label. After assigning functional labels, ChIP-modification emission and enrichment plots were reordered using ChromHMM Reorder for ease of visualization across species (**Fig. S1**). Finally, genomic intervals for each functional annotation were generated by merging all neighboring ChromHMM states with the same functional label.

ChromHMM regions assigned as ‘enhancer-like’ were only retained if they were flanked by ChromHMM regions assigned as unmarked.

### ChIP-seq signal quantification in regions of interest

For quantitation of ChIP-seq signal, aligned .bam files were converted to bedgraph files using deepTools bamCoverage (--normalizeUsing CPM --outFileFormat bedgraph), with the effective genome size parameter calculated using library read length and unique-kmers.py in khmer^75^. Read signal from bedgraph files was measured within regions of interest using BEDTools map (-c 4 -o sum -null 0 -F 0.5)^71^. The summed signal within intervals was averaged across biological replicates. To more clearly visualize the data, a Yeo-Johnson power transformation was applied. First, signal outliers were removed manually by visualizing the data and establishing appropriate signal thresholds, retaining an average of 99% of the original regions. Next, the transformation was performed using scipy.stats.yeojohnson^76,77^.

### Genomic characterization of ChromHMM regions

Various external genomic reference files were used to characterize features of our ChromHMM defined functional annotations:

#### Promoter regions

Promoters were defined as the two kilobase (kb) window surrounding transcriptional start sites present in Ensembl^78^ GTF files (human, rhesus, mouse, bull: Ensembl release 113; rat: Ensembl release 104), except for opossum where an Ensembl GTF was not available in monDom5 and a UCSC GTF^79^ was used. For some analyses, denoted in the figure legends, a single promoter window was chosen for each gene. Briefly, transcripts were prioritized using transcript support level when available (TSL=1), Ensembl “canonical tag” when available, or longest transcript when individual transcript confidence information was not available (rat, opossum).

#### GC content, CpG islands

GC content of individual regions was determined using BEDTools nuc and the relevant whole genome sequence file. Genomic coordinates for CpG islands were obtained via the UCSC Table browser (Group: Expression and Regulation; Track: CpG Islands) for all species.

#### Sequence conservation (phastCons)

Sequence conservation level was determined using mm39.phastCons35way.bw and hg38.phastCons100way.bw obtained from the UCSC Table Browser. phastCons bigwig tracks were converted to bedgraph using Kent utilities tool^80^ bigWigToBedgraph, and average signal within individual genomic regions was quantified using BEDTools map (-c 4 -o mean). For establishing a genomic phastCons average, bivalent regions in human and mouse were shuffled through the genome (BEDTools shuffle) and mean phastCons score was calculated in these regions.

#### Repeat elements

Repeat elements were obtained from UCSC Table browser (Group: Variation and repeats; Track: RepeatMasker) for all species, and repeats containing LINE, SINE, and LTR elements were retained.

### Gene ontology enrichment

GO enrichment was evaluated using the GOStats package^81^ in R, using all genes as the background set. *P* values were adjusted both by conditioning out child categories and by subsequent correction for multiple testing using the Benjamini–Hochberg method (p-adjusted). To determine the number of unique genes represented by selected GO terms (**Fig. S5F**), Amigo 2 (v.2.5.17)^82^ was used to download all human and mouse gene and gene products present within a GO term.

### De-novo motif analysis

STREME^37^ was used to identify enriched DNA-sequence motifs from bivalent regions of interest in human and mouse. For each region of interest, one kb of genomic sequence was obtained using the midpoint of the original region of interest. STREME was performed using either the default generated negative control sequences or a relevant set of sequences as the background control, as specified below. Each reported de-novo motif has an associated E-value, which is an estimate of the statistical significance of the motif compared to the negative control set.

Identified de-novo motifs were compared with known transcription factors (TF) binding motifs using Tomtom^83^. Three TF databases were used to compare to de-novo motifs identified: (JASPAR (2024^84^) core, non-redundant vertebrate motifs, HOCOMOCO (v11^85^) core, and human and mouse HT-SELEX motifs^86^). TFs were reported only if the de-novo motif and TF motif match had a q-value < 0.05 and if the TF was predicted in at least two databases.

To define enriched motifs across all bivalent regions in human and mouse, a random sample of bivalent regions nearby TSS (within +/- 1kb of TSS, n=10,000) was performed. De-novo motif enrichment analysis was performed with STREME using the default negative sequences. De-novo motifs were filtered by STREME E-value < 0.05 and the top 10 most frequent motifs (frequency defined as number of test sequences where the motif is present / total number of test sequences) were reported. To determine enriched motifs within evolutionary classes of bivalent regions, all human and mouse bivalent regions with the evolutionary labels ‘conserved’, ‘ancestrally active’, or ‘ancestrally Polycomb’ were used as the negative test set to distinguish sequences uniquely enriched for a given label compared to the other labels, and to exclude motifs generally enriched across all bivalent sites. Far fewer de-novo motifs were identified in this analysis; as such, up to ten of the most frequent motifs were shown. Position weight matrix graphs of identified motifs were generated using MEME suite tool ceqlogo.

### Defining spermatogenic and stable bivalent regions

To determine if bivalent regions in human and mouse germ cells were stable (also bivalent in ESC) or spermatogenic (not bivalent in ESC), we compared the ChromHMM defined bivalent regions between cell types. We performed a BEDTools intersect for all germ cell bivalent regions (-wao) compared to ESC bivalent regions. If a germ cell bivalent region had a minimum of 30% of the total size overlapping with an ESC bivalent region, the germ cell region was classified as “stable.” If a germ cell bivalent region had < 30% overlap with an ESC bivalent region, the germ cell region was classified as “spermatogenic.” All genomic intervals for “stable” and “spermatogenic” bivalent regions in human and mouse regions are provided in **Table S3**.

### Defining orthologous chromatin state between species

To determine orthologous chromatin states for ChromHMM-defined human and mouse elements in rhesus, rat, bull, and opossum, we first determined orthologous sequence elements in these species using liftOver^87^ (-minMatch 0.6), mapping regions from the human or mouse genome assembly onto the other genome assemblies. Regions which failed to map onto the second assembly were classified as “unmapped”. Once genomic regions were in the same assembly, we intersected the two species’ ChromHMM defined chromatin state regions using BEDTools intersect (-wao). If the human/mouse chromHMM region did not intersect with any chromHMM element from the other species, we categorized the chromatin state at the orthologous regions as “unmarked”. If the human/mouse region did intersect with a single element, we categorized the orthologous chromatin state as the functional annotation of that element as called in the other species. If the human/mouse region intersected with more than one element, we categorized the orthologous chromatin state as the functional annotation of the element called in the other species with the greatest amount of overlap (**Fig. S6A**). We performed this orthology method for each species to establish the orthologous chromatin state in each species for individual human/mouse regions.

### Determining evolutionary labels for bivalent regions

We next classified bivalent elements according to the orthologous chromatin state in other species. To define species specific bivalent regions, we retained regions where only the human/mouse region was bivalent, while no other orthologous chromatin states were allowed to be bivalent. Next, we “pruned” our phylogeny, assigning chromatin state within each lineage (ancestral, rodent, or primate) based on coherence between the two species in the group (human/rhesus, mouse/rat, or bull/opossum). For the ancestral group, we excluded orthologous regions where the chromatin state was discordant (i.e. an active region in bull and a bivalent region in opossum) but did not exclude regions where the chromatin state from one species was unknown (i.e. bivalent region in bull and an unmapped region in opossum), to account for lower rates of mapping in these species. The phylogeny of our chosen species allows analysis of two independent evolutionary trajectories: comparing the primate lineage vs. ancestral outgroups and comparing the rodent lineage vs. the same ancestral outgroups. For primate and rodent lineages, “conserved bivalent” regions were defined as those where both the ancestral species and the primate or rodent lineage were bivalent. “Ancestrally active” regions were defined as those where the ancestral group was classified as ‘activeTSS’ and the primate/rodent lineage was bivalent. “Ancestrally Polycomb” regions were defined as those where the ancestral chromatin state contained regions classified as Polycomb, allowing for some regions to be bivalent, and the primate/rodent lineage was bivalent. “Species specific” bivalent regions were defined as those regions where the orthologous chromatin state was mapped in at least one species within each group and all mapped orthologous chromatin states were not classified as bivalent (**Fig. S6B**). All genomic intervals for “conserved”, “ancestrally Polycomb”, “ancestrally active”, and “species specific” human and mouse regions are provided in **Table S5**.

### Determining evolutionary labels for genes

Genes were classified with an evolutionary label based on their promoter’s proximity to bivalent regions with an evolutionary label as described above. All gene promoter windows (2 kb) were intersected with bivalent elements classified as ‘ancestrally active’, ‘ancestrally polycomb’, ‘conserved’, or ‘species specific’ (BEDTools intersect -wao). Genes were then classified according to the evolutionary label of the bivalent region overlapping with the promoter window. Promoters overlapping with multiple bivalent regions with different evolutionary labels were excluded from analysis. The number of genes within each evolutionary category for human and mouse is shown in **Fig. 4A**. All genes classified as “conserved”, “ancestrally Polycomb”, “ancestrally active”, and “species specific” in human and mouse are provided in **Table S6**.

### Published datasets

All published data utilized in this study are listed in **Table S1**. All raw sequencing files were processed as described above. Details for further processing of data are provided here.

#### Bivalent promoters in other cell types

Mouse sperm H3K4me3 and H3K27me3 ChIP data was obtained from GSE42629^45^. Peaks were called using MACS3^88^ (-f BAM -g mm -q 0.10), using the sonicated genomic DNA as the control file (-c). For H3K27me3, broad peaks were called (--broad). Nearby peaks were merged using BEDTools merge (-d 500) and merged peaks with size >= 1kb were retained. To reduce false positive bivalent regions, H3K4me3 peaks were further filtered by MACS peak score (score >= 100). ChIP peaks from biological replicates were compared using BEDTools intersect and only regions with a peak in both replicates were retained. Finally, bivalent regions were defined by merging the filtered and replicated H3K4me3 and H3K27me3 regions (BEDTools merge -c 4 -o distinct) and regions were retained that had both an H3K4me3 and H3K27me3 peak (n=3,294). These bivalent regions were further filtered to retain those near promoters, requiring regions to overlap with a 2kb window centered on TSS (n=3,085).

GV oocyte bivalent regions were obtained from Hanna et al^17^. For bivalent promoters provided, a 2 kb window centered on the original midpoint of the region was utilized. For PGC^5^ bivalent genes, bivalent genes were required to be called as bivalent in at least one male PGC time point. For embryo (8-cell, morula, and ICM)^6^ and PGC bivalent promoters, a 2 kb window was defined at all TSS provided in the gene lists.

#### Somatic expression data and somatic expression breadth (tau)

Mouse somatic expression data (polyA+ RNA-seq, FPKM tables) were obtained from ENCODE, with all identifiers listed in Table S1. Processed human somatic expression data (TPM from RNASeQCv2.4.2) was obtained from GTEx^89^ (V10, accessed 10/21/2025). Briefly, tissue samples from male donors (Hardy Scale = 2) were considered. Using this criteria, expression data was obtained for different tissues, with variable numbers of individuals considered per tissue (brain (cortex): 88, brain (cerebellum): 92, heart (left ventricle): 75, adipose (subcutaneous): 110, kidney (cortex): 26, liver: 57, lung: 93, small intestine (terminal ileum): 8, spleen: 9, testis: 92, whole blood: 121). The gene expression level (TPM) was averaged across individuals for a given tissue type, and average TPM value is displayed. Tau, or expression breadth, was calculated for individual genes using expression values across all somatic tissue types with *tspex*^48^.

Processed bovine somatic expression data by tissue was obtained from CattleGTEx^52,90^ (Phase1 data). As in human, processed expression levels (TPM) were obtained within a tissue type, where the number of individual samples present within a tissue type varied (whole blood: 2195). The gene expression level (TPM) was averaged across individuals for a given tissue type, and average TPM value is displayed.

### Statistical methods

To compare the spatial co-occurrence of two sets of genomic regions (as in **Fig. 1J**), a Fisher’s exact test was used on the number of overlaps and unique intervals present within the regions sets using BEDTools fisher. To compare a distribution with a single control value (as in **Fig. 1G**), a 1-sample t-test was performed using scipy (ttest_1samp). To determine if two distributions were significantly different (as in **Fig. 2C**), a two sample Kolmogorov-Smirnov was performed using scipy (ks_2samp), To determine if two categorical variables were non-randomly associated (as in **Fig. 4G**), a Fisher’s exact test was performed using scipy fisher_exact. For all test that can be one or two sided, the test used is specified in the relevant figure legends.

### Visualization

ChIP-qPCR data was plotted using GraphPad Prism 10. All other data visualizations were generated in Python using Seaborn^91^ and Matplotlib^92^.

### Data access

All ChIP-seq and RNA-seq data generated for this project is available at GEO under accession number GSE326667. Reviewer access is available using token **ulknekmwxhizfav**. Accession numbers corresponding to public datasets used in this study are listed in **Tables S1** and **S2**.

## Acknowledgements

We thank K. Sumigray and A. Santos for help with rat tissue collection. We thank members of the Lesch laboratory for helpful discussions and assistance with protocols, in particular H. Yu and B. Walters. Pre-submission review was conducted using q.e.d science: qedscience.com.

Research reported in this publication was supported by the National Institute of General Medical Sciences of the National Institutes of Health under Award Number 1S10OD030363-01A1 to the Yale Center for Genome Analysis (YCGA). We are grateful to the YCGA for their sequencing contributions to this study. This work was supported by the National Institute of Child Health and Human Development (R01HD098128 and R21HD110843 to B.J.L.), and the National Cancer Institute (R21CA288677). Bluma Lesch, M.D., Ph.D. was supported by a Discovery Boost Grant, DBG-23-1150177-01-DMC, Grant DOI #10.53354/ACS.DBG-23-1150177-01-DMC.pc.gr.175431, from the American Cancer Society. B.J.L. is a Pew Scholar, supported by the Pew Charitable Trust.

## Author contributions

Conceptualization: DBF, BJL; Investigation: DBF, JT; Methodology: DBF; Formal analysis: DBF; Visualization: DBF; Writing – original draft: DBF; Writing – review & editing: BJL; Resources: BJL; Funding acquisition: BJL.

